# Identifying Promising Sequences For Protein Engineering Using A Deep Transformer Protein Language Model

**DOI:** 10.1101/2023.02.15.528697

**Authors:** Trevor S. Frisby, Christopher James Langmead

**Affiliations:** Computational Biology Department Carnegie Mellon University

**Keywords:** Protein design, Protein engineering, Attention, Transformer, Protein Language Model, Transfer learning, Fine-tuning

## Abstract

Protein engineers aim to discover and design novel sequences with targeted, desirable properties. Given the near limitless size of the protein sequence landscape, it is no surprise that these desirable sequences are often a relative rarity. This makes identifying such sequences a costly and time-consuming endeavor. In this work, we show how to use a deep Transformer Protein Language Model to identify sequences that have the most *promise*. Specifically, we use the model’s self-attention map to calculate a PROMISE SCORE that weights the relative importance of a given sequence according to predicted interactions with a specified binding partner. This PROMISE SCORE can then be used to identify strong binders worthy of further study and experimentation. We use the PROMISE SCORE within two protein engineering contexts— Nanobody (Nb) discovery and protein optimization. With Nb discovery, we show how the PROMISE SCORE provides an effective way to select lead sequences from Nb repertoires. With protein optimization, we show how to use the PROMISE SCORE to select site-specific mutagenesis experiments that identify a high percentage of improved sequences. In both cases, we also show how the self-attention map used to calculate the PROMISE SCORE can indicate which regions of a protein are involved in intermolecular interactions that drive the targeted property. Finally, we describe how to fine-tune the Transformer Protein Language Model to learn a predictive model for the targeted property, and discuss the capabilities and limitations of fine-tuning with and without knowledge transfer within the context of protein engineering.

## 1 Introduction

### 1.1 Protein engineering

Protein engineering is a rapidly evolving field that aims to develop novel protein sequences with useful properties. Protein engineers can be found across many scientific domains, so these useful properties can have equally far-reaching applications, including those that are therapeutic [1, 2, 3, 4], industrial [5, 6, 7], and environmental [8, 9]. Regardless of application, a common issue among protein engineering pipelines is the sheer number of experiments required to identify these useful sequences. The combinatorics alone associated with altering the amino acid composition at different sites within a protein sequence quickly escalates beyond what is experimentally feasible. Developing general-purpose methods that can help protein engineers with experimental decision making is thus important and necessary.

Protein engineering pipelines do not fit within a single mold. Just as the applications are varied, so too are the experimental requirements and focal points. While some workflows may focus specifically on a particular objective, many are multi-faceted and include different stages that each involve their own rounds of sequence selection or experimental decision making. A general workflow may be given as:

Protein Discovery → Protein Optimization → Model Building

As many protein engineering pipelines include at least one of these components, we will now briefly define each of these stages, and describe the types of experiments and decisions that are made within each.

#### Protein Discovery

While the design of novel protein sequences lies at the heart of most protein engineering objectives, in many settings there is an initial discovery phase. Perhaps most notably, this includes the design of antibody-based therapeutics [10, 11, 12]. Antibodies (Abs) play an important role in initiating an immune response by specifically targeting and binding targeted antigens. An Ab’s specificity for and ability to bind an antigen depends on the sequence identity at three complementarity-determining regions (CDR), as residues in these CDR will bind to the surface of an antigen. The Ab residues involved in antigen binding are known as paratopes, and those on the antigen as epitopes. Understanding the interactions between paratopes and epitopes is at the heart of many Ab therapeutics design problems [13, 14]. Recently, a similar class of proteins called nanobodies (Nbs) have also been used for their therapeutic properties [15, 16]. Nbs are shorter than Abs, which makes them easier and cheaper to produce, though are only found naturally in camelid species, which can make them harder to initially collect (whereas Abs can be obtained from common lab mice). Structurally, Abs and Nbs form Y-shaped proteins, where the CDR are located within hypervariable tips. The hypervariability means that sequences collected from an animal sample will have high diversity, and only a subset of the sequences collected will strongly bind a specified antigen target. Collecting this repertoire of diverse sequences is the crux of the protein discovery phase. These Abs or Nbs will undergo assays that quantify their ability to bind a specified antigen. From this, a panel of the best sequences, or *leads*, are identified and used in downstream engineering objectives that take advantage of the leads’ strong therapeutic properties. Finding ways to identify strong leads while minimizing the required wet-lab experimentation is thus an important undertaking.

#### Protein Optimization

Optimization is an integral component of all protein engineering campaigns. It is a stage where protein sequences are modified with the goal of identifying those that possess desirable properties. The optimization typically involves making a relatively small number of changes to a given parental protein sequence. For example, in *de novo* protein design [17, 18], computational models are used to propose novel sequences that are expected to have certain structural properties. These proposed sequences are typically experimentally refined (ie. optimized) to ensure they best match the desired properties. More commonly, naturally obtained sequences are optimized directly, such as leads selected from a protein discovery campaign. In any case, there are many types of experimental technologies that may be used to perform the sequence optimization. Traditionally, these approaches involve random mutagenesis at one or more selected sites [19]. Site-specific mutagenesis [20, 21, 22, 23] and deep-mutational scanning [24] further allow greater throughput and specificity as to where in a protein’s sequence edits should be made. Emerging approaches based on CRISPR/Cas9 [25, 26] continue this trend. Given the costs associated with these experiments as well as the exponential number of *potential* edits that can be made to any protein sequence, it is important to identify ways that efficiently and effectively select at which sites to perform these mutagenesis experiments.

#### Model Building

In conjunction with the novel sequences that may be designed and synthesized during a protein engineering campaign, it is often desirable to also obtain a structural model of the system under investigation. The model serves as a tool that can be used to better understand physical properties of a protein, such as how it folds or interacts with other proteins. Traditionally, these models have been obtained through experimental means, such as x-ray crystallography [27]. While x-ray crystallography can provide a high-resolution structure of a protein or protein complex, it requires very specific conditions that may preclude its applicability to certain systems. It also only provides a static, or fixed model for a protein’s structure, whereas most interactions worth modeling depend on many moving parts. *Computational* models built on machine learning and artificial intelligence bridge this gap between the generalizability and applicability of purely experimental model building approaches. With respect to protein structure prediction, AlphaFold [28] is able to predict any protein’s 3D structure, and is shown to have highest accuracy on the benchmark CASP challenges [29]. More generally, computational models can be used to make predictions on any protein property. In some settings, these models can even form a feedback loop, where predictions by a model are used to select experiments, and the experimental results are used to update the model [30, 31, 32]. Whether or not a model is used in-line with running protein engineering experiments, most protein engineering pipelines generate large amounts of data, which makes them well-suited for training machine learning models at the conclusion of all experimentation.

### 1.2 Representation learning and Transformer Protein Language Models

Protein engineers look to design novel proteins that have certain desired functions or structure. It is has been shown that these biological features are represented through the statistical dependencies between evolutionarily selected sequences that are found in nature [33, 34]. Finding ways to encode and exploit this evolutionary information could help to guide the sequence design choices made by protein engineers. Traditionally, these statistical dependencies have been conveyed through multiple sequence alignments (MSA). MSAs are used to infer patterns that exist between protein sequences, such as those that are conserved or co-vary, both of which can influence protein structure and function [35]. A machine learning technique known as *generative modeling* can allow one to sample new protein sequences that maintain the underlying statistics of a given MSA [36]. However, the computational complexity of sequence alignment can make it difficult to perform across large data sets, and most of the millions of publicly available protein sequences are unaligned and unlabeled [37].

The development of high-capacity generative language models that are trained on unlabeled data has been an area of keen focus within the field of Natural Language Processing (NLP). The Transformer [38, 39] is one such class of model that has been successfully applied to many NLP tasks [40, 41]. While the composition of protein sequences (ie. the 20 natural amino acids) is considerably different than text used in NLP tasks, recent work has shown that the Transformer has great promise when applied to protein sequences [42]. The authors show that, after training on over 250 million distinct protein sequences, their Transformer model, which they call the Evolutionary Scale Model (ESM-1b), successfully encodes biochemical properties of amino acids, biological variation and remote homology, and secondary structure and tertiary contacts.

ESM-1b, as well as other Transformers generally, works by learning a *representation* for an input sequence. This representation is a high dimensional, real-valued vector that encodes the statistical dependencies and patterns present in the sequences used during training. With ESM-1b, this representation is learned by means of a deep neural network. This network consists of 33 connected Transformer blocks, each of which consist of an attention mechanism that feeds into a feed-forward neural network. The attention mechanism [43] is what allows the model to capture the statistical dependencies across all sequences, as it accounts for all pairwise interactions between each position in a sequence. The resulting self-attention map, or matrix of all pairwise interactions, can thus be interpreted as a protein-protein interaction map.

Since ESM-1b is trained on many millions of protein sequences spanning practically all known protein families, it learns many general patterns that exist across all protein sequences. When model building, it is typical to want to learn how to predict a specific task (eg. how proteins from a specific family binds a specific target). Transformer models can provide a good starting point when obtaining these specific models. Transformers have been shown to be effective *few-shot learners* [44], meaning they can be used to predict specific tasks when given only few labeled examples. Developing exact mechanisms to train such models is an active area of research within NLP, and includes methods such as knowledge distillation [45], fine-tuning [46], and transfer learning [47].

In this work, we show how to use the deep transformer protein language model ESM-1b to help select experiments associated with the protein discovery and protein optimization phases of protein engineering pipelines. We then demonstrate how to apply the computational model building techniques of fine-tuning and transfer learning using the Transformer in each of these settings.

## 2 Materials and Methods

### 2.1 Encoding a sequence and its protein target as input for ESM-1b

We use the deep Transformer Protein Language Model ESM-1b [42] to jointly model interactions between a protein sequence, *s_binder_*, and its protein binding target sequence, *s_target_* (Figure 1). For our purposes, ESM-1b is a function ESM(·) that takes as input a length *n* (tokenized) protein sequence, *s*, and outputs a tuple 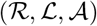:

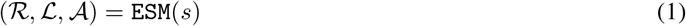

**Figure 1:**
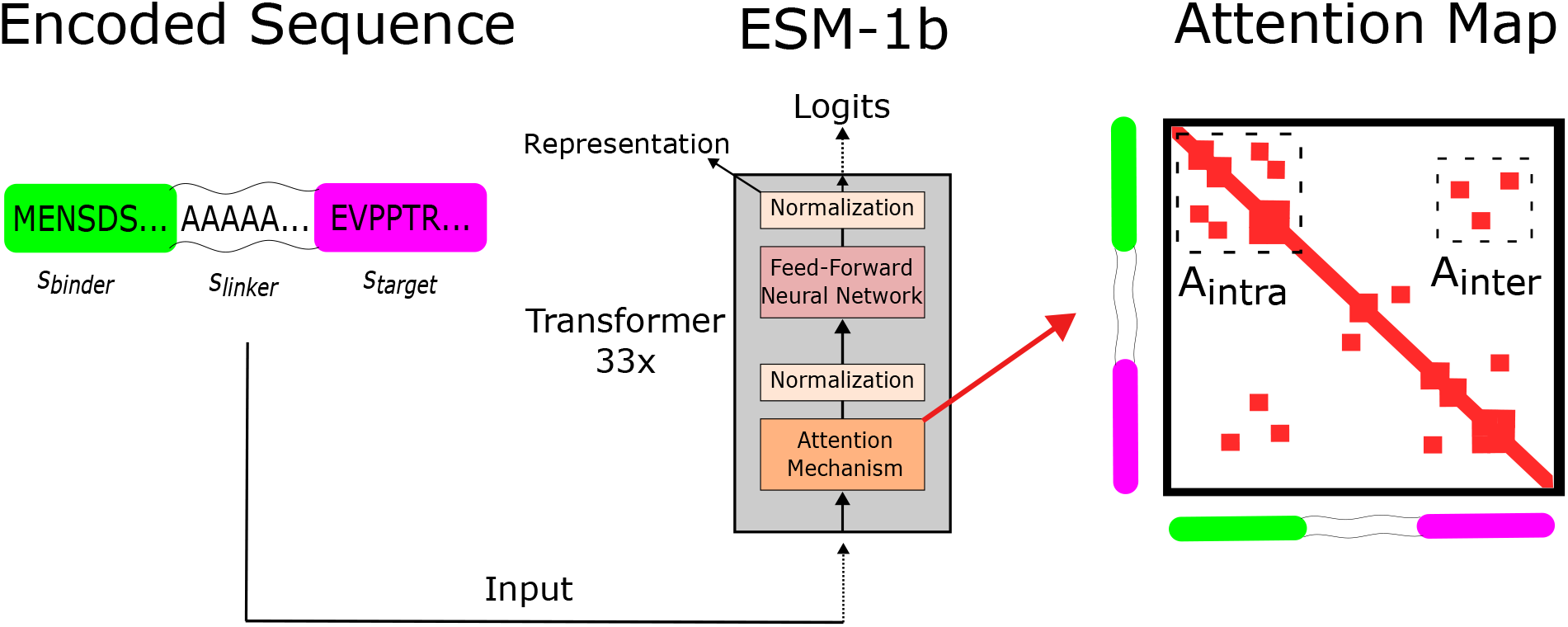
We use a deep Transformer Protein Language Model to identify *promising* sequences within protein engineering campaigns. Using Nb discovery and protein optimization as two test cases, we introduce the Promise score, and show how it can help address the needle-in-the-haystack issue of identifying protein sequences with targeted properties. To calculate a Promise score, we specifically encode a sequence and its binding target through concatenation with a linker, and use this adjoined sequence as input to the pre-trained ESM-1b model. This Transformer model uses an attention mechanism that captures all expected pairwise interactions between residues within the input sequence through an attention map. We use this map to quantify intermolecular and intramolecular interactions within each protein sequence. The Promise score reflects these interactions, meaning stronger sequences should tend to have higher Promise score than weaker sequences.

Here, 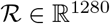 is the sequence *representation*, or feature-rich learned encoding obtained from the final Transformer block. 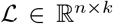 contains the *logits* returned by the Transformer model. These logits can be used to assign an evolutionary probability to each of *k* possible tokens (the 20 standard amino acids, nonstandard amino acids, and special tokens internal to ESM-1b) at each position of the input sequence. Finally,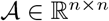 is the self-attention map returned by the final Transformer block’s attention mechanism. This attention map is analogous to a predicted protein-protein interaction map, and is the principal output from ESM-1b that we use.

We want to use ESM-1b to identify interactions between *s_binder_* and *s_target_*. To do this, we construct input sequence *s* to encode both of these sequences jointly. In our experiments, we show that a simple concatenation scheme is effective. We encode each input sequence *s* as:

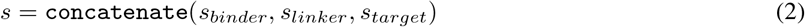

where *s_linker_* is a sequence of one of the following:

1. **Alanine**. An amino acid. A poly-alanine linker represents a flexible loop in *s*.
2. **<mask>**. A special token used by ESM-1b.
3. **No linker**. *s_binder_* and *s_target_* are concatenated without a separating linker.

ESM-1b imposes the constraint that any input protein sequence must be no longer than 1024 residues. In our experiments, the combined lengths of *s_binder_*, *s_linker_* and *s_target_* was always considerably less than 1024. We experimented with linker lengths 10, 50, and 100.

### 2.2 **Formulating a** “Promise score”

In order to assess the promise of a given sequence *s_binder_* as a focus for protein engineering, we use the attention map for *s* to calculate a “Promise score”. Treating the attention map as a predicted protein-protein interaction map, we use it to calculate a score that accounts for *intermolecular* interactions between *s_binder_* and *s_target_*, and, when appropriate, *intramolecular* interactions within *s_binder_*. Let *n*, *ℓ*, and *τ* be the lengths of *s_binder_*, *s_linker_*, and *s_target_*, respectfully, and let *N* = *n* + *ℓ* + *τ*. We accomplish this by first isolating sections of that correspond to either intermolecular or intramolecular interactions:

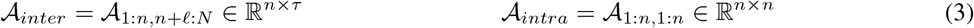

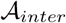 and 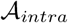 represent intermolecular and intramolecular interaction profiles, which we use to calculate separate intermolecular and intramolecular scores. We obtain these profiles and scores for all sequences being considered for protein engineering. To standardize the numerical scaling of these profiles in order to ensure fair comparisons between different sequences, we apply min-max scaling to both as follows:

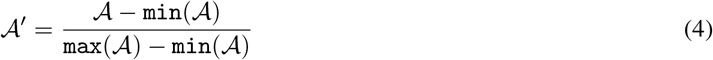

The intermolecular interaction score *S_inter_* is given as a sum over 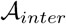 normalized by the length of *s_binder_*. For all 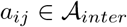:

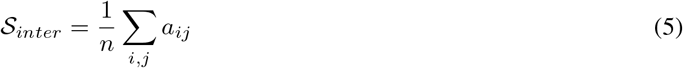

In our protein discovery experiments, we simply use Promise score 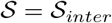. In many real-life protein engineering pipelines, a subsequent protein optimization stage applies site-specific mutagenesis experiments to a functional but suboptimal baseline protein sequence *s_baseline_*. This could correspond to a lead sequence selected from a previous protein discovery campaign, or a wildtype sequence that has been identified previously. We obtain intramolecular interaction score 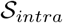 by calculating a difference between the intramolecular interaction profiles of *s_baseline_* and *s_binder_*. Where 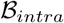 is the intramolecular interaction profile for *s_baseline_*, we obtain such a score as:

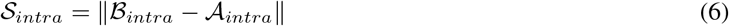

The larger 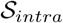 is, the greater the difference between intramolecular interaction profiles. We assume that the functional property of *s_baseline_* is driven in part by the the intramolecular interactions reflected by 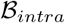. This leads us to expect that a functional 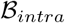 should have an intramolecular interaction profile that is more similar than dissimilar to 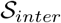. This reasoning motivates our formulation of the Promise score during protein optimization— a linear combination of 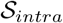 and 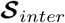:

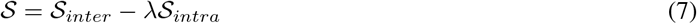

A promising design will have many intermolecular interactions and an intramolecular interaction profile that is similar to a known functional baseline.

Hyperparameter *λ* ≥ 0 governs the degree to which the intramolecular term impacts the overall score. In principle, any standard hyperparameter fitting method could be applied to set *λ*. We have devised a means to automatically select *λ* that takes advantage of the fact that *S_inter_* and *S_intra_* can be pre-computed for any sequence without the need for an external label. Let 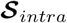 and 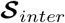 be the set of all 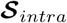 and 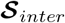 computed for each sequence that is being evaluated. Our approach is to treat *λ* as a scaling factor between 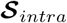 and 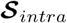. Let 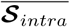 refer to the mean of 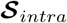, and 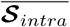 the mean of 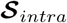. We identify the scaling factor by taking a simple ratio, and use this approach to select *λ* in all of our experiments:

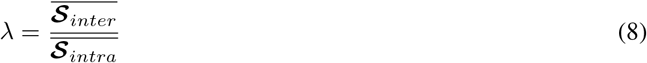

### 2.3 Fine-tuning and knowledge transfer with ESM-1b

While we have so far used the pre-trained ESM-1b model as-is in order to calculate Promise Scores, we now describe how it can be *fine-tuned* in order to learn a model that predicts the binding strength between *s_binder_* and *s_target_*. In order to imbue ESM-1b with the ability to provide such a prediction, we attach a three-layered neural network to the head of the model. In other words, ESM-1b’s learned representation serves as input for the neural network head. Fine-tuning occurs during model training. During backpropogation, we fine-tune by updating the parameters of ESM-1b’s final Transformer block in conjuction with the parameters of the neural network head. All other ESM-1b parameters are left fixed at their pre-trained state. In our experiments, the neural network head consists of three linear layers connected by rectified linear unit (ReLU) activation functions. The full model is trained for 30 epochs using PyTorch’s AdamW optimizer [48, 49] with default parameter settings (learning rate *γ* = 1e−6, *β*_1_ = 0.9, *β*_2_ = 0.999, and weight decay coefficient *λ* = 0.01).

We expect that it would be beneficial to be able to take knowledge gained in one engineering campaign and apply it to that of another (related) campaign. To this end, we show the ability to transfer knowledge learned from one protein engineering objective to another within the discovery and optimization stages. Within protein discovery, we use models for protein binding learned from Nano-HSA data and show their ability to predict binding on Nano-GST data, and vice-versa. Similarly within protein optimization, we take models learned from BRCA1-BARD1 binding data and show their ability to predict binding between Spike and ACE2, and vice-versa.

### 2.4 Data

We use the Promise score at two phases of the protein engineering pipeline— protein discovery and protein optimization. Table 1 provides an overview of the data we use in our experiments.

**Table 1:**
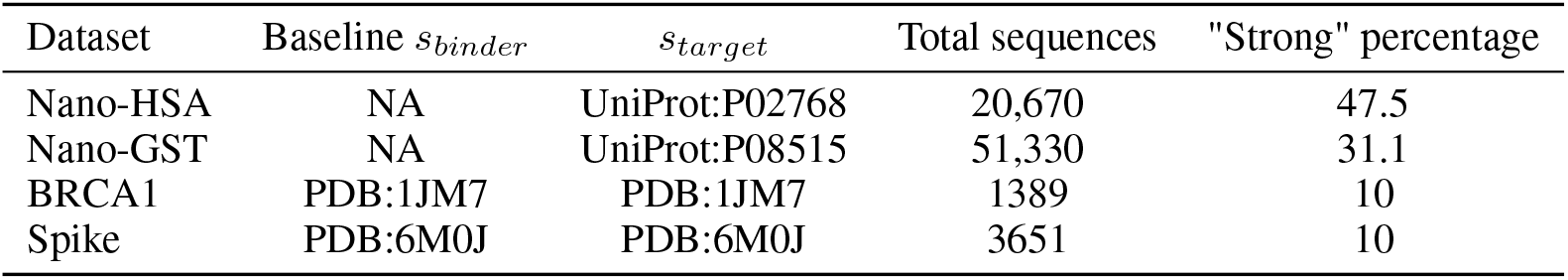
An overview of the data we used to evaluate the Promise score. The sequence IDs are given as the DataBase:ID indicating where the sequence can be accessed, or are NA (not applicable).

At the protein discovery phase, we show how to use the Promise score to select lead sequences from two Nb repertoire data sets [50]. The repertoires were previously collected from an immunized llama using protein targets human serum albumin (Nano-HSA) and glutathione S-transferase (Nano-GST). The relative binding strengths of each Nb sequence to its target sequence was originally provided as a categorical label, where one of the three possible labels corresponds to strong binding sequences. We re-labeled each sequence as “strong binding” or “not strong binding” to obtain a binary descriptor for relative binding strength, which we use in our experiments. In total, the Nano-HSA data set contains 20,670 Nb sequences and the Nano-GST data set 51,330. The percentage of strong binders in Nano-HSA and Nano-GST was 47.5% and 31.1%, respectively.

When computing Promise score 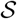 with these data, we let *s_binder_* be the full Nb sequence. However, when obtaining 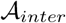, we focused our attention on the CDR3 region of each Nb by only considering the region of attention map 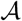 that corresponds to intermolecular interactions between the CDR3 and target sequence.

At the protein optimization stage, we show how to use the Promise score to select mutagenesis experiments for two different proteins— BRCA1 and Spike. We obtained BRCA1 sequences from single site mutational scans [51] that measured the downstream activity caused by interaction between BRCA1 and its binding target protein BARD1. These measurements were obtained from 1389 single site mutations. Spike sequences were similarly obtained from single site mutational scans [52] that measured the binding affinity between Spike and its binding target protein ACE2, and it included 3651 single site mutations. We obtained binary labels for both datasets by binning the raw experimental values, where the top 10% were labeled as “strong” sequences, and the rest “not strong”.

## 3 Experiments and Results

### 3.1 The Promise Scores **of strong and weak binders are different**

In order for the Promise score to provide an effective means for selecting sequences for protein engineering, the scores of known strong binders should be distinguishable from those of weak binders. According to how the Promise score is formulated, we further expect that the scores of strong binders should be generally higher than those of weak binders. For both Nb discovery and protein optimization, we computed Promise score for all sequences using different linker types and lengths, and compare how strong binders were scored relative to weak ones. The raw scores are only comparable within a given linker type and length— the relative ability to discern strong from weak binders is what can be compared.

During Nb discovery (Figure 2 top), there is a significant difference in the distribution of Promise score of strong and weak binders for each linker type and length, and the average score of strong scoring sequences is higher than that of weak binders. For Nano-GST, using no linker obtains Promise score that have greatest ability to discern strong from weak binding Nb sequences on average. For Nano-HSA, a length 100 alanine linker provides the greatest ability to discern. For both Nano-HSA and Nano-GST, calculating a Promise score using a mask linker of any length or a length 50 alanine also confers the ability to discern strong from weak binders (Figure S1 top).

**Figure 2:**
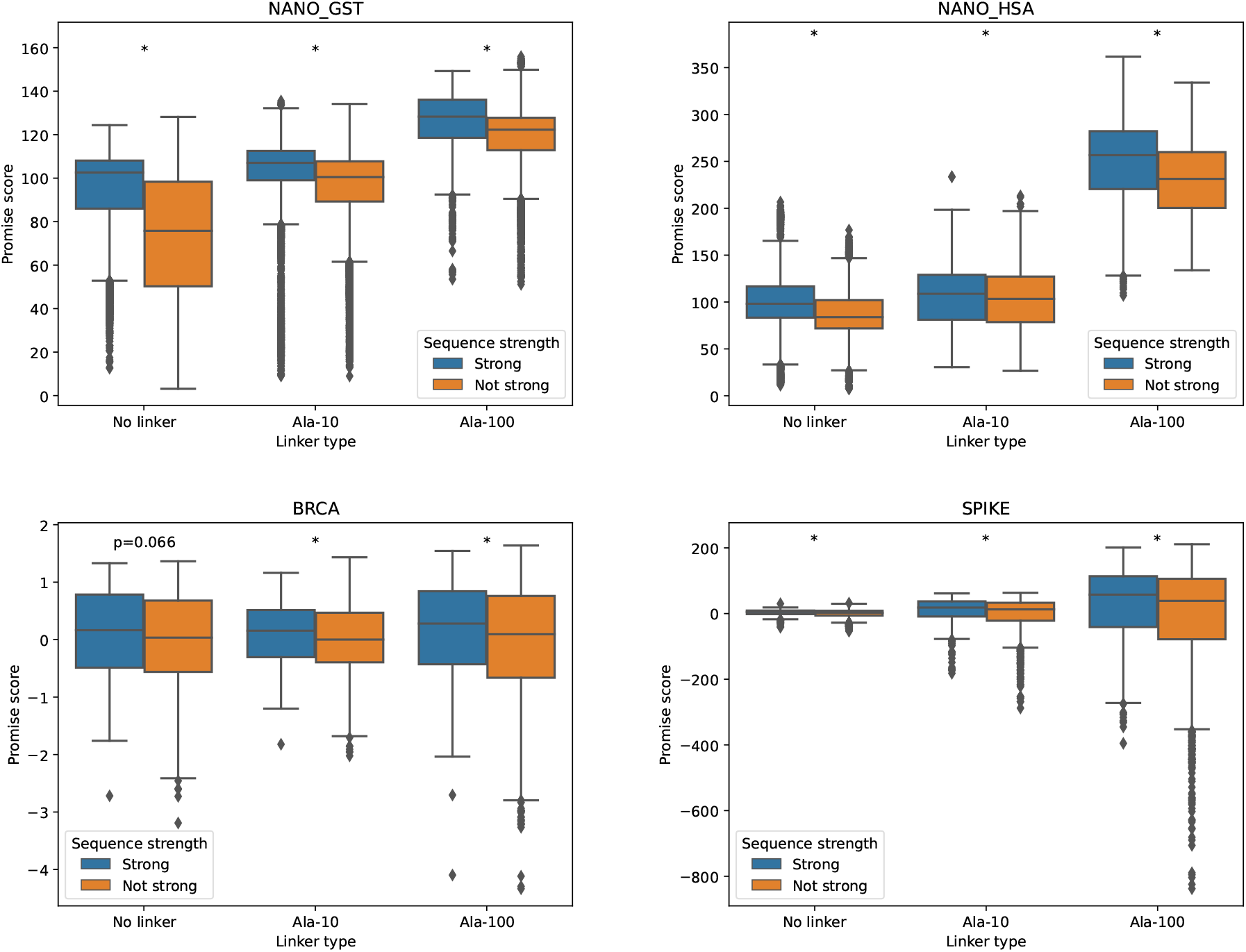
We calculated Promise score for each sequence identified within two Nb discovery campaigns (Top) and two protein optimization campaigns (Bottom), and compare how “strong” sequences scored compared to those that were “not strong”. In each case, strong binders had an average Promise score greater than sequences that were not strong. These differences were either statistically significant (*p*< 0.05, Mann-Whitney U-test, denoted with *), or had a p-value just above the 0.05 threshold (BRCA no linker– *p* = 0.066).

During protein optimization (Figure 2 bottom), in all but two cases, there is again a significant difference in the distribution of Promise score of strong and weak binders for each linker type and length, where the stronger sequences have a higher Promise score on average than their weak counterparts. The exceptions are with BRCA1 using no linker and a length 10 mask linker, though they just barely misses the threshold for significance at *p* = 0.066 and *p* = 0.054, respectively. In general, the Promise score for each linker type and length in protein discovery and protein optimization provides a narrow (but still real) ability to discern strong from weak binders. This also includes using a mask linker or length 50 alanine (Figure S1 bottom).

### 3.2 The Promise score **identifies beneficial experiments**

Since strong binders tend to have higher Promise score than weak binders, we can use the score to rank the sequences in a repertoire and expect that a set of top-ranked sequences will be enriched with strong binders. Figure 3 demonstrates this for different sized subsets obtained by taking the top 1, 2*,…*, 100 sequences ranked by the Promise score. Thus, the Promise score may be useful as a means to identify binders.

**Figure 3:**
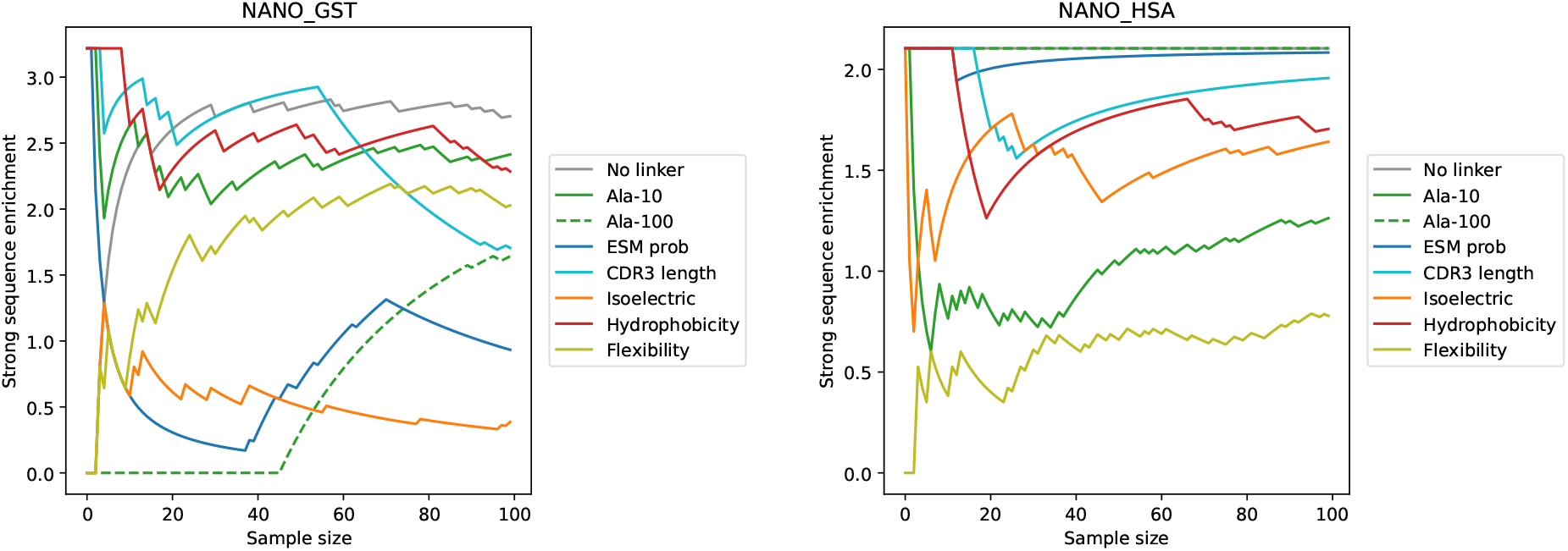
We used Promise Scores to help identify potential lead sequences from two Nb discovery campaigns. We found that Promise score derived methods identified sequences that were enriched with strong binders, and often identified a higher frequency of strong binders compared to other sequence-based calculation methods. In both Nb discovery campaigns, the best overall strategy used the Promise score.

During protein discovery, we compare the cumulative strong binder enrichment for the top 100 sequences obtained using the Promise score calculated with no linker, 10 length alanine, and 100 length alanine linkers. The enrichment is given as the ratio of the observed strong binder frequency at a given sample size to the expected strong binder frequency (ie. the underlying true strong binder frequency). We compare to sequence-derived calculations commonly used to assess Nb sequences when identifying potential lead sequences. Though none of these metrics directly measure a protein’s ability to bind a target, they measure qualities that are reasonably expected to correlate with this property. These metrics include:

1. **CDR3 Length**. The number of residues within the CDR3. While an optimal CDR3 length depends on the particular target sequence, previous analysis [50] of the Nb sequences used in our experiments determined that sequences with shorter CDR3 tended to be stronger binders.
2. **Nb Isoelectric point (pI)**. [53, 54] The pH where the Nb carries no net electrical charge. Low pI Nbs are more likely to aggregate, which can lead to inactivation and induces immunogenicity [55].
3. **CDR3 Hydrophobicity.** The frequency of hydrophobic residues within the CDR3. The prevalence of hydrophobic residues can increase the propensity for aggregation [56].
4. **Nb Flexibility**. [57] The relative mobility of Nb residues. An Nb with greater flexibility may have greater ability to bind its target.

We ranked all sequences according to these metrics and report the cumulative strong binder enrichment of the top 100 sequences. CDR3 length and hydrophobicity are ranked in ascending order, whereas Nb isoelectric point and flexibility are ranked in descending order. We also used the logits produced by ESM-1b to calculate a probability under the ESM-1b model for the CDR3 of each sequence. We do this by computing the softmax over the logits associated with each position of the CDR3. The probability is then given as the product over the probabilities assigned to each residue present in the sequence. We only used the Nb sequences as input for ESM-1b when calculating a probability, rather than the sequence concatenated to the target sequence, and report the cumulative strong binder enrichment of the top 100 most probable sequences.

With Nano-GST (Figure 3 left), the Promise score computed with no linker and a length 10 alanine linker have a cumulative strong binder enrichment greater than two at almost every sample size up to 100. At sample size 100, they produce two of the highest cumulative strong binder enrichment values, along with a length 50 alanine linker (Figure S2 left). Using the Promise score computed with a length 100 alanine linker struggles to identify strong binders for the top 50 scoring sequences, but by sample size 100 produces a cumulative enrichment greater than 1.5. Calculating a Promise score using a mask linker tends to have a strong binder enrichment less than 2 (Figure S2 left). Among the sequence-based calculations, hydrophobicity tends to produce the highest cumulative enrichment, and typically ranks between no linker and alanine 10. The shortest CDR3 are enriched with strong binders, but the enrichment fades by sample size 100. The isoelectric point of Nb sequences is generally not enriched for strong binders at these sample sizes, and neither are the probable sequences under the ESM-1b model.

With Nano-HSA (Figure 3 right), the Promise score computed with no linker and a length 100 alanine linker are consistently the two strategies that yield the greatest strong binder enrichment. Both have cumulative enrichment greater than two across all sample sizes up to 100. Using any length mask linker or 50 length alanine yields an enrichment greater than 1.5 for each sample size up to 100 (Figure S2 right). The most probable sequences under the ESM-1b model also fare well, with a strong binder enrichment consistently right around 2. The Promise score calculated with a length 10 alanine linker has the lowest strong binder enrichment among the ESM-1b derived metrics. Each of CDR3 length, CDR3 hydrophobicity, and Nb isoelectric points tend to have cumulative strong binder enrichment that is lower than using no linker or a length 100 linker, but higher than the length 10 linker. Sequences predicted to have high flexibility are not enriched for strong binders.

To summarize across both Nb discovery campaigns, the no linker Promise score most consistently identified sequences enriched with strong binders. Among the comparison metrics, each of the isoelectric point, protein flexibility, and ESM-1b were inconsistent– sequences selected according to each were enriched with strong binders in one campaign and not in the other. While both hydrophobicity and CDR3 length consistently identified sequences enriched with strong binders, at most sample sizes, the enrichment was lower than that obtained by the no linker Promise score.

We also show how the Promise score can be used to select site-specific mutagenesis experiments during protein optimization. Rather than selecting from a set of discovered sequences, the task here is to select specific positions from a baseline sequence at which to conduct single-site mutagenesis. The intent is to select the positions that will cumulatively yield the highest percentage of strong binders. To do this, we first obtain the Promise score for each individual (linked) variant sequence. We then take the top 500 scoring sequences, and identify at which position each sequence is variant relative to the baseline sequence. We treat this tally as a “vote", and the selected sites correspond to the positions that obtain the most votes. In our experiments, we identified the top 10 sites. Figure 4 shows the strong binder frequency obtained by this procedure on the BRCA1 and Spike protein optimization sequences.

**Figure 4:**
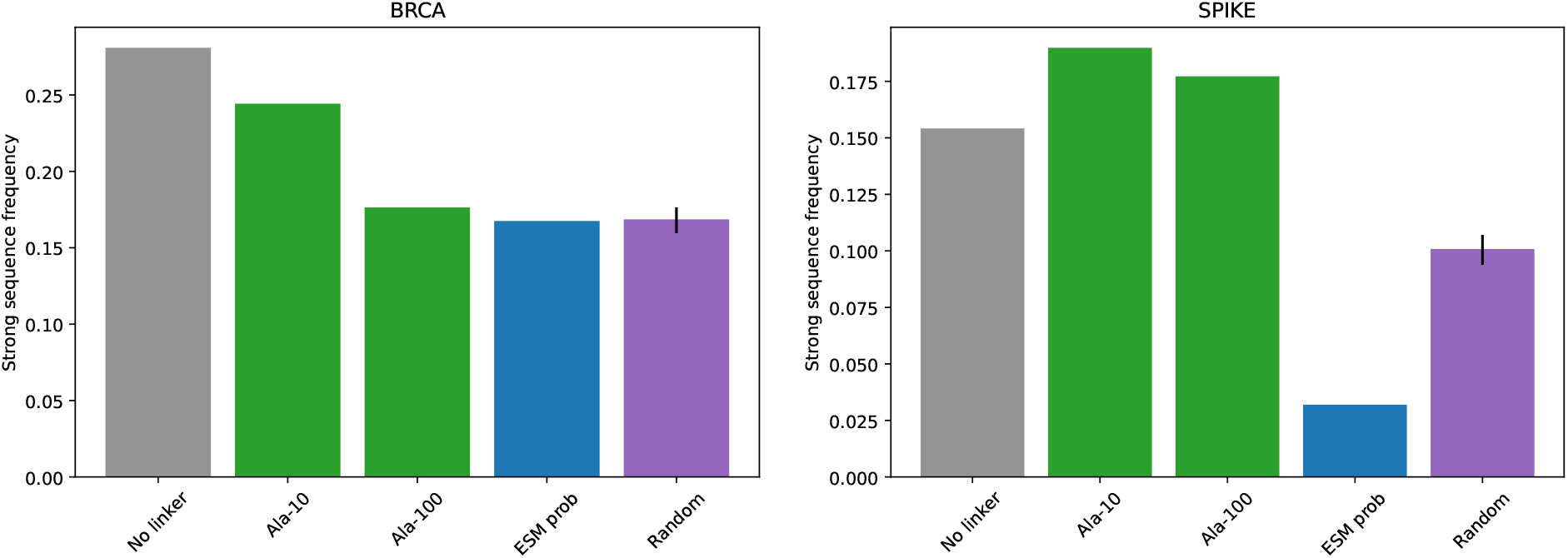
We used Promise score to select 10 site-specific mutagenesis experiments on each of BRCA1 and Spike. For most linker types and lengths, the Promise Scores identified sites that had a higher strong binder frequency than selecting randomly or using evolutionary sequence probabilities.

With BRCA1 (Figure 2 left), the Promise score calculated with no linker identified sites that yielded the highest percentage of strong binders. Alanine with length 10 linker also performed well and had the second highest strong binder frequency. The length 100 alanine linker had the third highest strong binder frequency. We generally found that short mask linkers were also effective (Figure S3 left). To better quantify how enriched for strong binders these experiments were, we simulated randomly selecting mutagenesis sites 50 times for comparison. Both no linker and a length 10 alanine linker have greater strong frequency enrichment relative to the random sampling. The length 100 alanine linker performed comparably to random sampling. We also calculated the probability of each variant sequence under the ESM-1b model, and used those probabilities to select single sites for mutagenesis. We used the same voting scheme as with the Promise score, but instead used the sequence probabilities. The sites with most probable sequences yielded strong binders at frequency similar to random sampling.

With Spike (Figure 2 right), the Promise score calculated with alanine linkers yield the highest strong binder frequency. The length 10 linker was highest and the length 100 linker second highest. Using no linker yielded the third highest frequency. All three of these approaches performed similarly to each other and better than random sampling. We found that using a mask linker was generally less effective than using an alanine linker (Figure S3 right). The sites selected by the most probable sequences under the ESM-1b model were not enriched for strong binders, and had a lower strong binder frequency than random sampling.

### 3.3 The attention map provides insights into protein-target interactions

In addition to identifying strong binders for the purposes of protein engineering, Promise score can also provide biological insights about the modes of interaction between a protein sequence and its target protein. To calculate any Promise score, we use the attention map to quantify intermolecular interactions between a protein sequence *s_binder_* and its target *s_target_*, given by 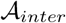. The rows of 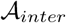 correspond to each position of *s_binder_*, whereas the columns correspond to each position of *s_target_*. While calculating the Promise score involves summing over all of 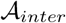, we can sum over the columns to instead obtain a per-residue score for the target sequence. We use these per-residue scores to identify specific residues on *s_target_* that are expected to interact the most with a given *s_binder_*. To identify the target residues predicted to be most involved in intermolecular interactions, we took the top 10% of protein sequences according to Promise score, and used these sequences to obtain an average per-residue score for the target residue. We then identified target residues that scored in the top 20 percentile as those most strongly believed to be involved in intermolecular interactions. Figure 5 shows 3D structures of each target protein tested with the top scoring residues highlighted (Figure S4 shows each of these structures rotated 180^*°*^).

**Figure 5:**
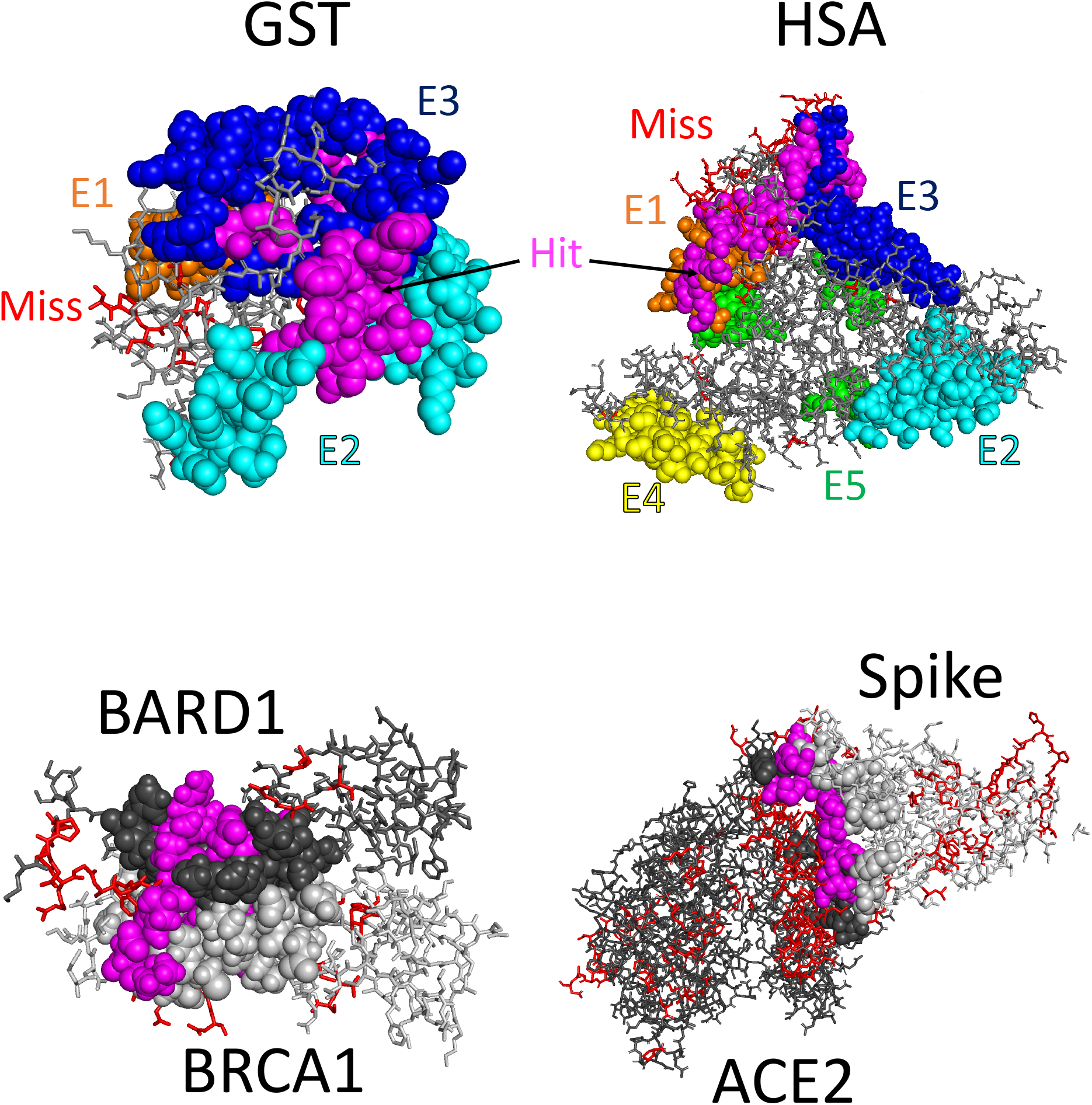
We use intermolecular contact map 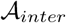 to calculate residue-specific scores to identify residues that contribute the most to intermolecular interactions. With Nb discovery (Top), we highlight overlap between our top scoring residues and epitopes previously validated through cross-linking mass spectrometry. The magenta spheres indicate this overlap. Each other colored sphere indicates a residue within a validated epitope region. The red sticks correspond to top scoring sites that did not fall in any validated region (denoted “Miss”), and the gray sticks indicate all other residues within the protein sequence. With protein optimization (Bottom), we show structures of each protein bound to its target. The spheres indicate residues known to play a role in protein-target binding. Those shaded magenta were identified by the per-residue scoring. Red sticks indicate residues outside this binding region that were also identified. Dark gray sticks indicate *s_target_* residues, and light gray *s_binder_* residues. The structures we used are given by PDB IDs 1DUG, 1AO6, 1JM7, and 6M0J.

In a Nb discovery campaign, identifying target residues involved in intermolecular interaction is akin to identifying Nb epitopes. With both Nano-GST and Nano-HSA, we compared the strongest scoring residues on target proteins to experimentally validated epitope regions. With Nano-GST (Figure 5 top left), we identified 42 GST residues predicted to be involved in intermolecular interactions. Of these identified residues, 12 of them are located within the experimentally validated E3 epitope. E3 was the strongest validated epitope, as 50% of the tested Nbs bound to this region. Seven of the 12 residues are singletons within the E3 region located at residues 158-200, and the other five residues are at the contiguous E3 region at residues 213-217. Of the remaining identified residues, one fell within the E2 epitope region (residue 125), and the rest did not fall within one of the validated epitope regions.

With Nano-HSA (Figure 5 top right), we identified 98 HSA residues predicted to be involved in intermolecular interactions. These predicted interacting residues overlap most strongly with validated epitopes E1 and E3. For E1, these include the contiguous region at residues 23-25, 113-114, and 126-127, as well as the singleton residues 28, 111, 116, 118, 123, and 138. For E3, these include the contiguous regions at residues 5-13 and 93-94 as well as singletons at residues 96 and 98. 5% of the validated Nbs bound to epitope E1 and 20% at epitope E3. There were two sporadic singleton residues identified in other validated epitopes, residue 172 in E5, and residue 301 in E2. The remaining selected residues did not fall in one of the validated epitope regions, though blanketed the protein region surrounding epitopes E1, E3, and E5.

We performed similar analyses with both of the protein optimization protein sequences. In addition to computing per-residue scores for *s_target_*, we also computed per-residue scores for *s_binder_* by summing over the rows of 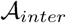. As with the *s_target_* per-residue scores, we identified the top 20% of residue scores as those most involved in intermolecular interactions. We found 3D structures for both the BRCA1-BARD1 (Figure 5 bottom left) and Spike-ACE2 complexes (Figure 5 bottom right), and highlight the residues on *s_binder_* and *s_target_* that had highest scores indicative of inter-molecular interactions. In general, we found that the largest contiguous cluster of identified residues were located at the protein-target binding interface.

Binding between BRCA1 and BARD1 is known to be driven by interactions between BRCA1 residues 8-22 and 81-96 and BARD1 residues 36-48 and 101-116 [58]. Of the 23 high scoring BRCA1 residues we identified, 10 of them fell within these regions (residues 14-15, 18-19, 21-22, 86, 93, and 95-96), and the remaining high scoring residues flanked each of these regions (residues 24, 26-27, 41-42, 79, and 97-103). Similarly, 10 out of the 24 BARD1 residues we identified fell within these ranges (residues 36-38, 42, 45, 102, 105, 108-109, and 112), and the rest also flanked these regions (residues 27-35, 49, 51, 54, 99, and 134).

Spike-ACE2 binding is facilitated through specific interactions between 17 different Spike residues and 20 different ACE2 residues [59]. We identified 39 Spike residues expected to be most involved in intermolecular interaction, two of which overlap with the known interacting residues. While some of the non-overlapping residues are in the vicinity of these binding regions, we found that identified residues were widely spread across the entire Spike protein. With ACE2, we identified 121 residues, including 13 of the known interacting residues. While many of the non-overlapping residues were again in the vicinity of the known binding region, many were also spread throughout the ACE2 protein.

In order to demonstrate that the per-residue patterns we have described above did not arise by chance, we simulated *randomly* selecting residues for each protein system. In each case, we randomly selected a number of residues equal to the amount identified by the per-residue Promise score. Over 50 replicates, we identified the frequency with which these randomly selected residues fell within a known binding area, and compared these frequencies to those obtained with the Promise score selected residues (Figure S5). Indeed, for each of the regions that we identified substantial overlap between the residues selected by the per-residue Promise score and a known binding region, we found that the Promise score selected residues within that region at a rate higher than if selected at random.

### 3.4 Using ESM-1b to learn models for protein binding

Having used ESM-1b to prioritize sequences expected to strongly bind their target, we now show to what extent we can use ESM-1b to learn predictive classification models for this targeted property. With Nb discovery, we randomly selected 5000 sequences from Nano-GST and Nano-HSA to serve as two separate training sets, and another 15,000 sequences from each as testing sets. Within a protein engineering context, the training sequences are analogous to a preliminary round of experimentation carried out for the purpose of collecting initial data. We then used fine-tuning to train two classifiers, one that was trained with Nano-GST, and one that was trained with Nano-HSA. We then used these models in a traditional (ie. used to predict binding strength to the same target sequence used during training) and transfer learning manner. As a point for comparison, we also trained classifiers without fine-tuning. That is, during training, we only updated the parameters of the neural network head, and left all ESM-1b parameters fixed. This demonstrates the ability to simply use the pre-trained representations when learning a model for protein binding. We also used these models in both the transfer and traditional learning settings.

We obtained comparable results with both Nano-GST and Nano-HSA (Figure 6 top). Fine-tuned models used traditionally were far and away the best at predicting binding strength. The model used to predict Nb binding strength with GST had an AUC of 0.988, and the model used to predict Nb binding strength to HSA an AUC of 0.958. The non-fine-tuned models that used pre-trained representations were second best, though clearly less effective than their fine-tuned counterparts. The AUC obtained were 0.795 with GST and 0.753 with HSA. We found that knowledge transfer hurt the predictive capability of both the fine-tuned and non-fine-tuned models. The non-fine-tuned models were hurt less, and still achieved an AUC of 0.712 with GST and 0.683 with HSA. Fine tuning combined with transfer learning essentially yielded random classifiers, as the models has AUC of 0.514 and 0.474 with GST and HSA, respectively. We found similar results for each linker type used in the input sequence encoding (Figure S6).

**Figure 6:**
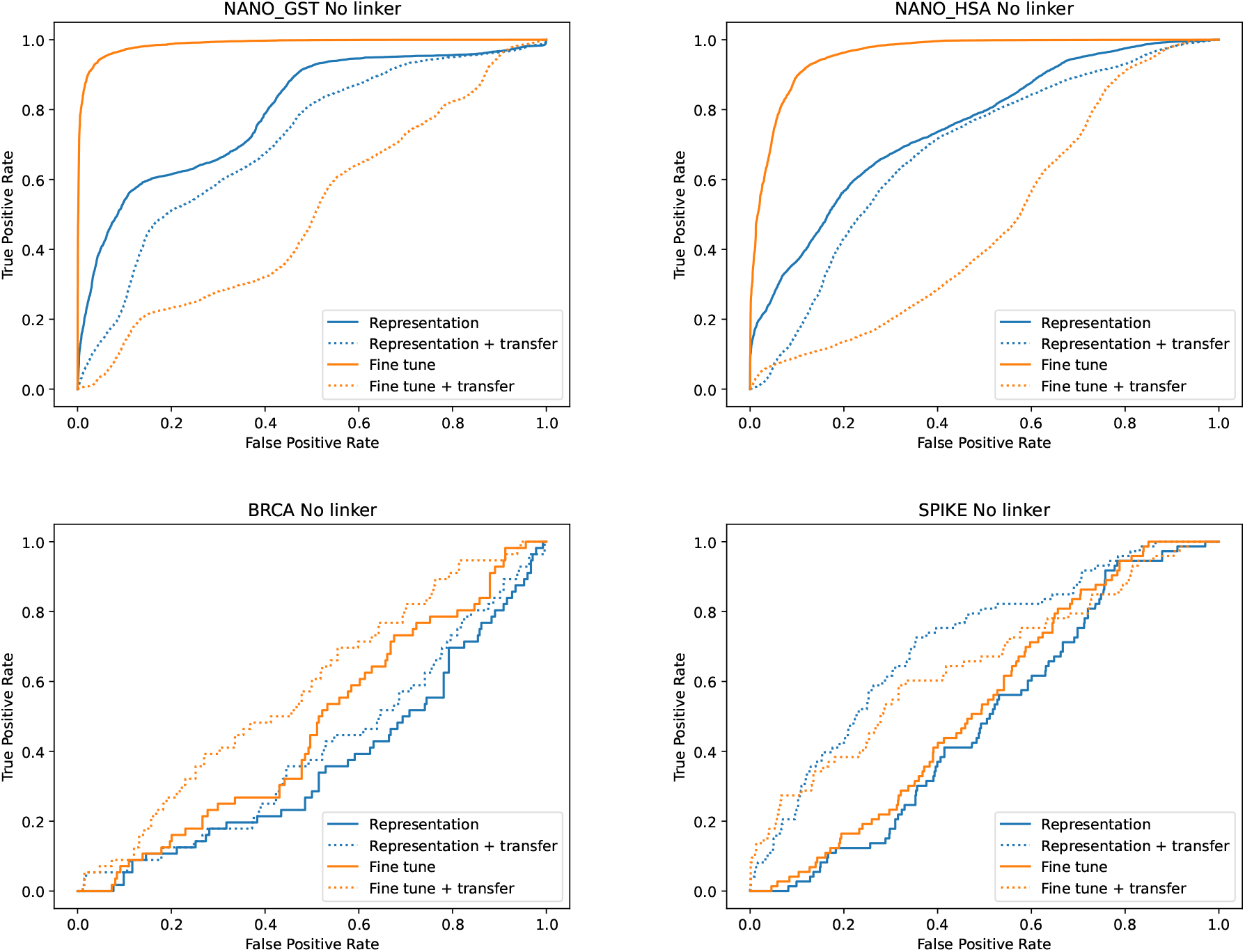
ROC curves for models trained to classify strong and weak binders for both Nb discovery (top) and protein optimization (bottom). Orange curves correspond to models that used fine-tuning, and blue those that did not. Solid curves correspond to models applied in a traditional machine learning paradigm, whereas dashed lines used transfer learning. With Nb discovery, fine-tuned models used in the traditional setting were very strong. While all models learned in protein optimization were of lower quality, we saw evidence that transfer learner led to model improvement.

We next used ESM-1b to train classifiers using the two sets of protein optimization sequences. For both BRCA1 and Spike, we randomly selected 75% of the single residue sites in each protein. We used the sequences previously obtained via single-site mutagenesis at these selected positions as training data. The mutatations at all other sites were then used as test data. We again trained models with and without fine-tuning, and used both in a traditional and transfer setting.

While we were able to obtain at least one reliable classification model for protein binding using fine-tuning with the Nb discovery data, we found that it was much more difficult to do so within the context of protein optimization (Figure 6 bottom). With BRCA, the best model used fine-tuning with transfer, but only had an AUC of 0.573, meaning the model does not effectively classify strong and weak binders. Interestingly, the ROC curves provide some evidence that transfer learning may have some positive effect relative to traditional learning, though again none of the models score highly according to AUC. With Spike, we again found that the models that used transfer learning outperformed those that used traditional learning. We obtained the best model using no fine-tuning with transfer, which had an AUC of 0.708. The model with fine-tuning plus transfer had an AUC of 0.646. The traditional learning models performed no better than random classifiers. As with Nb discovery, we found that using different linkers had minimal effect on the resulting models (Figure S7).

## 4 Discussion

We have shown how to use the deep Transformer Protein Language Model ESM-1b to calculate a Promise score that can prioritize specific sequences and be used to guide protein engineers towards designs that are more likely to bind to a given target. We believe this can lead to more efficient and successful protein engineering campaigns.

### 4.1 **The role of linkers when computing** Promise score

In our experiments, we hypothesized that self-self attention maps would reveal, in a coarse-grained fashion, the interactions between a sequence *s_binder_*, and a binding partner *s_target_*. We tested this hypothesis by first concatenating *s_binder_* and *s_target_* with a separating linker, and use this as input to the ESM-1b model. Strictly speaking, ESM-1b was not designed with this use case in mind. Rather, the model was trained using millions of *monomer* sequences, and has been used to describe secondary structures or make downstream predictions for *singular* protein sequences. And yet, when we use the attention map generated with our concatenated protein-target sequence, we are able to calculate Promise score that tend to favor strong binders. From a biological perspective, this suggests that the types of interactions present in the monomers used to train ESM-1b are informative enough to identify the intermolecular interaction between a protein and its binding partner. From a modeling perspective, this suggests that representation learning approaches trained on monomers are strong enough to identify the ways that multiple sequences interact with each other.

Where we used linkers, we experimented with two types — a polyalanine, and the special <mask> character that is internal to ESM-1b. Alanine’s simple methyl side chain makes it a flexible amino acid. The idea was that the model may interpret a chain of alanines as a “tether” connecting the protein sequence and its target protein sequence. The flexibility would allow the two protein sequences to interact as if they were two unconnected proteins in close proximity. The <mask> character is used when training ESM-1b. Random residues are replaced with the <mask> character, and the model learns to predict which residue belongs where the <mask> is placed. In our use, the <mask> character denotes space between sequences without specifically committing to any particular residue type that may bias the interactions between residues.

Generally speaking, we found that using no linker always produced a good result. We found that using a linker could occasionally perform better than no linker, but not overwhelmingly so. When comparing the two linker types, we found that the natural amino acid alanine produced more consistent results, whereas there was much greater variance in the performance of Promise score calculated with <mask> linkers. For future work, we would recommend using no linker or a short alanine linker.

### 4.2 **The generalizability of** Promise score

We first showed how to use Promise score to select lead sequences using Nbs obtained from two different Nb discovery campaigns. Notably, we showed that Promise score selected sequences are often stronger than sequences selected by a number of sequence-based scoring metrics. This again shows the power of representation-learning approaches like ESM-1b. Despite not being trained to know anything about the biophysical properties of a given protein sequence, we show that we can use the model to make inferences about the relative strength of a protein with greater effect than using biophysical calculations.

We also showed how to use Promise score to select single site mutagenesis experiments during protein optimization. The primary task associated with protein optimization is considerably different than that of Nb discovery. Rather than identifying a subset of promising sequences from a naturally large and diverse repertoire, the goal is to identify specific sites on a baseline sequence to perform single-site saturation mutagenesis. As a result, the set of sequences under consideration will be smaller and more uniform than those encountered during Nb discovery, both of which add to the challenge of this task. We overcame this challenge by adding an extra term 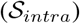 to the calculation of Promise score 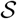 that accounted for intramolecular interactions. The formulation of 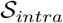 is consistent with protein optimization— we want to identify sequences that enhance an already existing feature of a baseline sequence. So while we have shown the effectiveness of computing particular Promise score for Nb discovery and protein optimization, the more general point is that we have shown an ability to formulate scoring functions using the attention map of a Transformer Protein Language Model that are tailored to specific protein engineering objectives. We believe that our work can serve as a baseline for different protein engineering objectives that have different constraints and assumptions.

While we have demonstrated how the Promise score may be used in isolation to prioritize protein sequences during Nb discovery and protein optimization, we know that most protein engineering pipelines are multi-facteted. They often include different types of analyses that have somewhat orthogonal yet complementary focuses. These may include genomic assays that identify DNA-protein interactions [60], molecular modeling that predict molecular dynamics and structure [61], or phylogenetic analyses that uncover genomic relationships between related species [62]. We want to emphasize that using Promise score is similarly complementary, as they can be used to prioritize sequences at any and all stages of complex protein engineering pipelines.

Finally, we believe it is important to note that Promise score can be computed using any Transformer Protein Language Model, not just ESM-1b. While we found ESM-1b to be an effective and convenient model with which to showcase this ability, any model that uses self-attention could be used in its place. Our intent is not to champion one Transformer model over the other, but to demonstrate that this class of model can be used to prioritize protein sequences within the context of protein engineering. We see this as a strength of the approach, as it can be easy to implement using new and improved models and they become available. Indeed, even the author’s of ESM-1b have released other versions of Transformer Protein Language Models with differing designs and specifications since we began working with ESM-1b [63, 64, 65]. We leave formal comparisons between Transformer Protein Language Models applied towards this goal to future work.

### 4.3 ESM-1b as a coarse-grained model for intermolecular interactions

While the Promise score is a singular numeric value that reflects the promise of a sequence in its entirety, we showed that we can also easily obtain per-residue scores for *s_binder_* or *s_target_*. Using these per-residue scores, we show that we can identify specific residues that are known to be involved in intermolecular interactions. More generally, we believe these results demonstrate that ESM-1b can be viewed as a coarse-grained model for intermolecular interactions. With each set of sequences that we used, we observed that the “hits” and “misses” tended to congregate in regions of each protein that have highest concentration of known interacting residues.

In the case of Nb discovery, we used these per-residue scores to identify potential epitopes, and compared these findings to experimentally validated epitope sites. It is important to note that the experimental validation consisted of cross-linking models at sites chosen by computational molecular docking. The validation thus does not provide any evidence for or against potential epitopes beyond the sites that were tested. Additionally, the experimental validation only used 24 Nb sequences, meaning the reported binding percentages are noisy estimates of the true underlying values. Still, across two sets of Nb discovery sequences, the per-residue scores identified sites that overlapped three out of eight total validated epitope regions, as well as identifying many sites outside these regions. We believe that this is an encouraging sign that Protein Language Models like ESM-1b are able to identify specific residues that are expected to contribute to intermolecular interactions. Such an ability could allow protein engineers to better pinpoint the regions in a protein of interest that should be investigated within an protein optimization campaign. Of course, we also made simplifying assumptions in our work that contribute to disagreement with the validation analyses. For one, we only considered interactions involving CDR3, so any effect caused by CDR1 or CDR2 are unaccounted for entirely. Additionally, we arbitrarily used the top 20 percentile sequences to calculate the per-residue scores, where conceivably there is some other optimal threshold. As such, we see these analyses as a jumping off point for future work that identify epitope, or even paratope sequences.

Applied to protein optimization, we investigated not only finding per-residue scores for *s_target_*, but also doing so for *s_binder_*. For the BRCA1-BARD1 complex, nearly half of the residues that we identified fell within the known binding interface. This strongly suggests that high per-residue scores obtained from 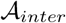 do indeed identify residues involved in intermolecular interactions. In the case of Spike-ACE2, we identified more than half of the known interacting ACE2 residues, though only two out of the 17 known interacting Spike residues. This highlights an important distinction behind how we compute scores for each *s_target_* (ACE2) and *s_binder_* (Spike). For each sequence used, *s_target_* is unchanged, whereas each *s_binder_* differs from all others at exactly one residue (due to each Spike sequence having been obtained from a single site mutation scan). Thus, the distribution of high scores throughout the Spike sequence indicates that the model believes different mutations affect the residues that interact at the binding interface. This capability could be used to generate hypotheses about how mutations affect binding patterns between a protein and its partner. Still, that the vast majority of known ACE2 interacting residues were identified indicates that the Transformer accurately identifies interacting residues within the Spike-ACE2 complex.

### 4.4 Model learning within the context of protein engineering

Having a predictive model for the targeted property in a protein engineering campaign could serve as an invaluable asset, as it can serve as a tool that helps with sequence design decisions. For this reason, we investigated how well we could learn such models using ESM-1b. First, we compared fine-tuning ESM-1b to using the sequence representations from the pre-trained model, and found that fine-tuning worked very well with the Nb discovery setting. With protein optimization, we found it difficult to learn reliable models with or without fine-tuning. We believe this ability to learn reliable models with Nb discovery but not with protein optimization can be explained by differences between these data. As discussed in section 4.2, the Nb discovery data is both more diverse and contains more sequences than the protein optimization data. Diversity and data quantity are well-known factors that influence the ability to train machine learning models [66]. This highlights an important consideration for the experimental design choices within protein engineering campaigns. Our work suggests that single-site mutagenesis experiments across relatively short spans of a protein sequence may not yield informative enough data for accurate model learning. If obtaining a reliable predictive model is imperative or desirable when using single-site mutagenesis, protein engineers may want to consider ways to inject greater diversity into their experiments.

In addition to fine-tuning, we also investigated the viability of knowledge transfer when training models. The ability to use knowledge transfer would allow protein engineers to take what they learn during one engineering campaign and apply it to another. With the Nb discovery data, we found that knowledge transfer was less harmful compared to not using knowledge transfer if using the pre-trained representations rather than a fine-tuned model. This makes sense— a fine-tuned model is adapted to work with a specific *s_binder_*-*s_target_* pair, so using it on different data would be counter productive. Interestingly, with the protein optimization data, we found that using knowledge transfer yielded models with higher AUC relative to using no knowledge transfer. One thought as to why we may see an improvement in these cases again relates back to the issue of diversity. Perhaps being trained on a set of sequences that were different entirely than those being tested on actually provided a greater degree of diversity that led to slightly better performance. As a final point, we only tried a very basic form of transfer learning in each of these cases, where we use one data set to train a model (eg. Nano-HSA), and make prediction on a separate data set (eg. Nano-GST). It is conceivable that having access to many different data sets to use during training could improve the model’s performance.

## 5 Conclusion

We have shown how to use the deep Transformer Protein Language Model ESM-1b to identify promising sequences during protein engineering campaigns. By jointly encoding a protein sequence *s_binder_* and its binding target *s_target_*, we use ESM-1b’s self-attention mechanism to identify intermolecular and intramolecular interactions between these sequences. We use these interactions to formulate what we refer to as the Promise score, and show how this score can be tailored to prioritize protein sequences in two distinct protein engineering domains— protein discovery and protein optimization. With protein discovery, we show how to use Promise score to effectively select lead sequences from two separate Nb repertoires. And with protein optimization, we show how to use Promise score to identify single-site mutagenesis experiments that successfully identify strong binders. In both cases, we show how to also compute per-residue scores that indicate those expected to undergo intermolecular interactions. We showcase that high scoring residues on Nb target proteins correspond to known epitopes, and those within the BRCA1-BARD1 and Spike-ACE2 complexes correspond to known interacting residues. Finally, we demonstrate that ESM-1b can be fine-tuned to learn accurate models for Nb binding strength, and discuss the limitations of model learning in single-site mutagenesis protein engineering campaigns. We believe this work is highly adaptable and directly applicable to many protein engineering pipelines, so can help to make protein engineering more efficient and effective.

## Acknowledgments

We would like to thank AMD for the donation of critical hardware and support resources from its HPC Fund that made this work possible.

