## Supplemental Material for "Identifying Promising Sequences For Protein Engineering Using A Deep Transformer Protein Language Model"

February 15, 2023

### 1 Supplemental Results

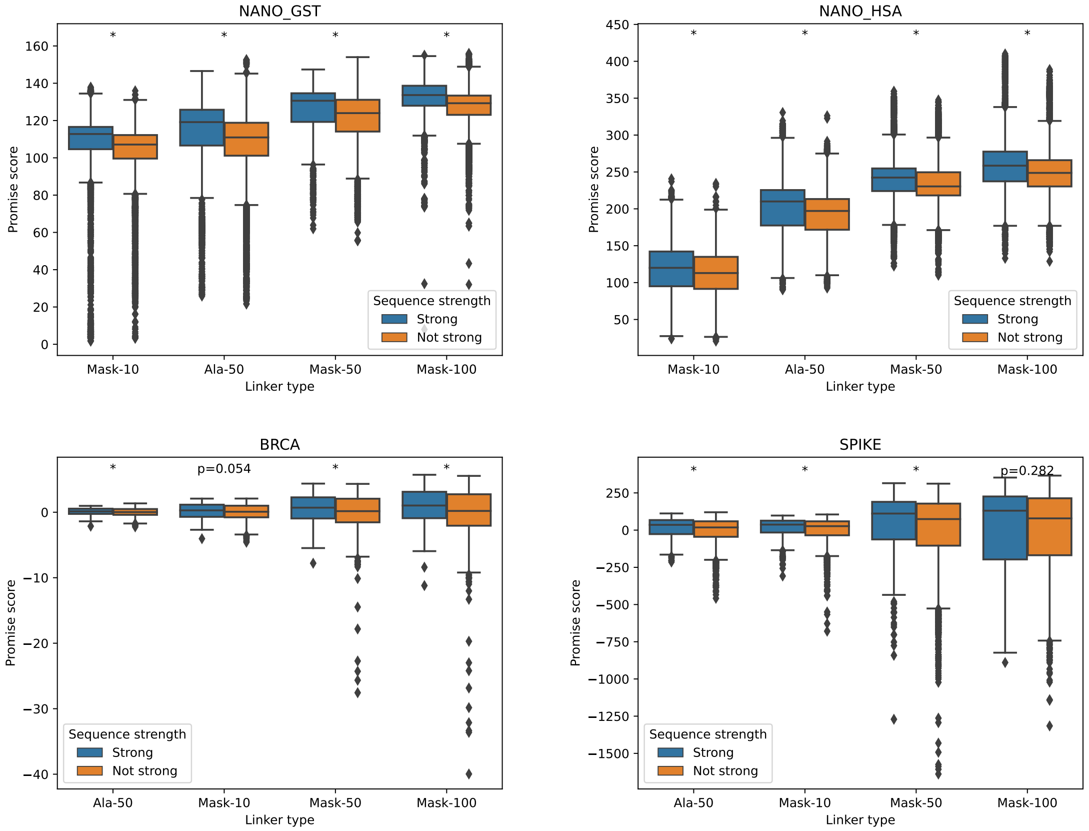


Figure S1: We calculated PROMISE SCORES for each sequence identified within two Nb discovery campaigns (Top) and two protein optimization campaigns (Bottom), and compare how “strong” sequences scored compared to those that were “not strong”. In each case, strong sequences had an average PROMISE SCORE greater than sequences that were not strong. These differences were either statistically significant ($p < 0.05$, Mann-Whitney U-test, denoted with *) or had a p-value just above the 0.05 threshold (BRCA mask-10– $p = 0.054$), with the only exception when using a 100 length mask linker with Spike ($p = 0.282$).

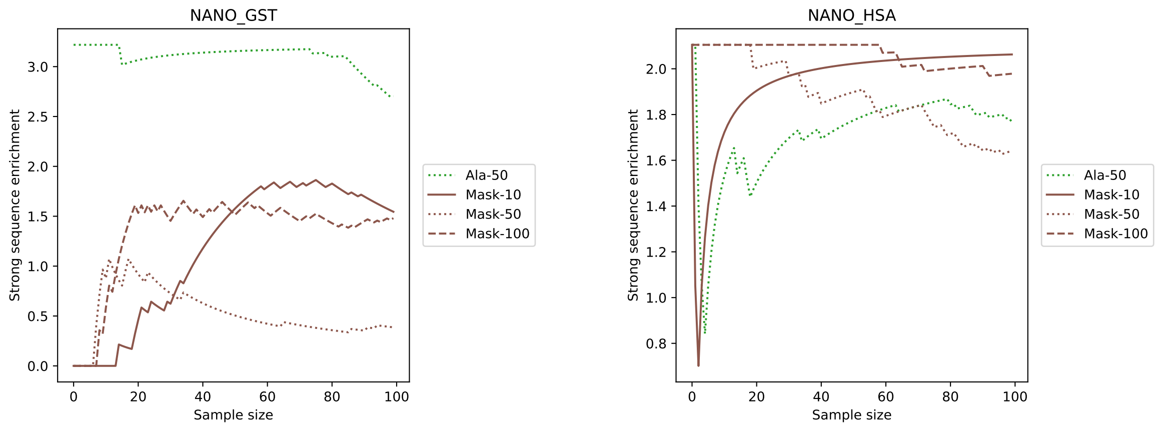


Figure S2: We used PROMISE SCORES to help identify potential lead sequences from two Nb discovery campaigns. Here, we show the strong sequence enrichment when using PROMISE SCORES obtained with mask linkers of length 10, 50, and 100, as well as a length 50 alanine linker. With Nano-GST, we found that alanine linkers yielded higher strong sequence enrichment than mask linkers. With Nano-HSA, we found that all linker types yielded comparable strong sequence enrichments.

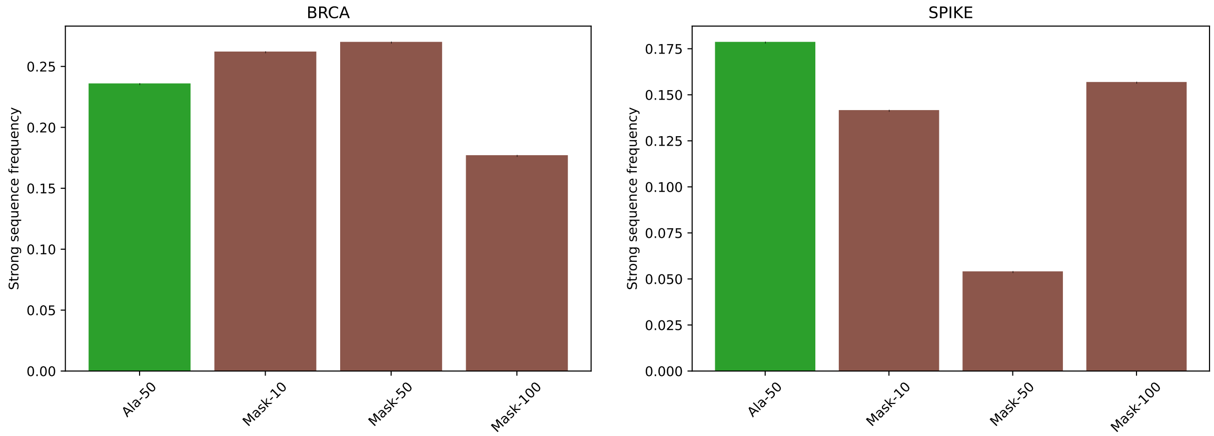


Figure S3: We used PROMISE SCORES to select 10 site-specific mutagenesis experiments on each of BRCA1 and Spike. With BRCA1, using length 10 or 50 mask linkers yield strong sequence frequencies comparable to those when using alanine linkers. With Spike, the mask linker yields more variable results compared to alanine linkers at different linker lengths.

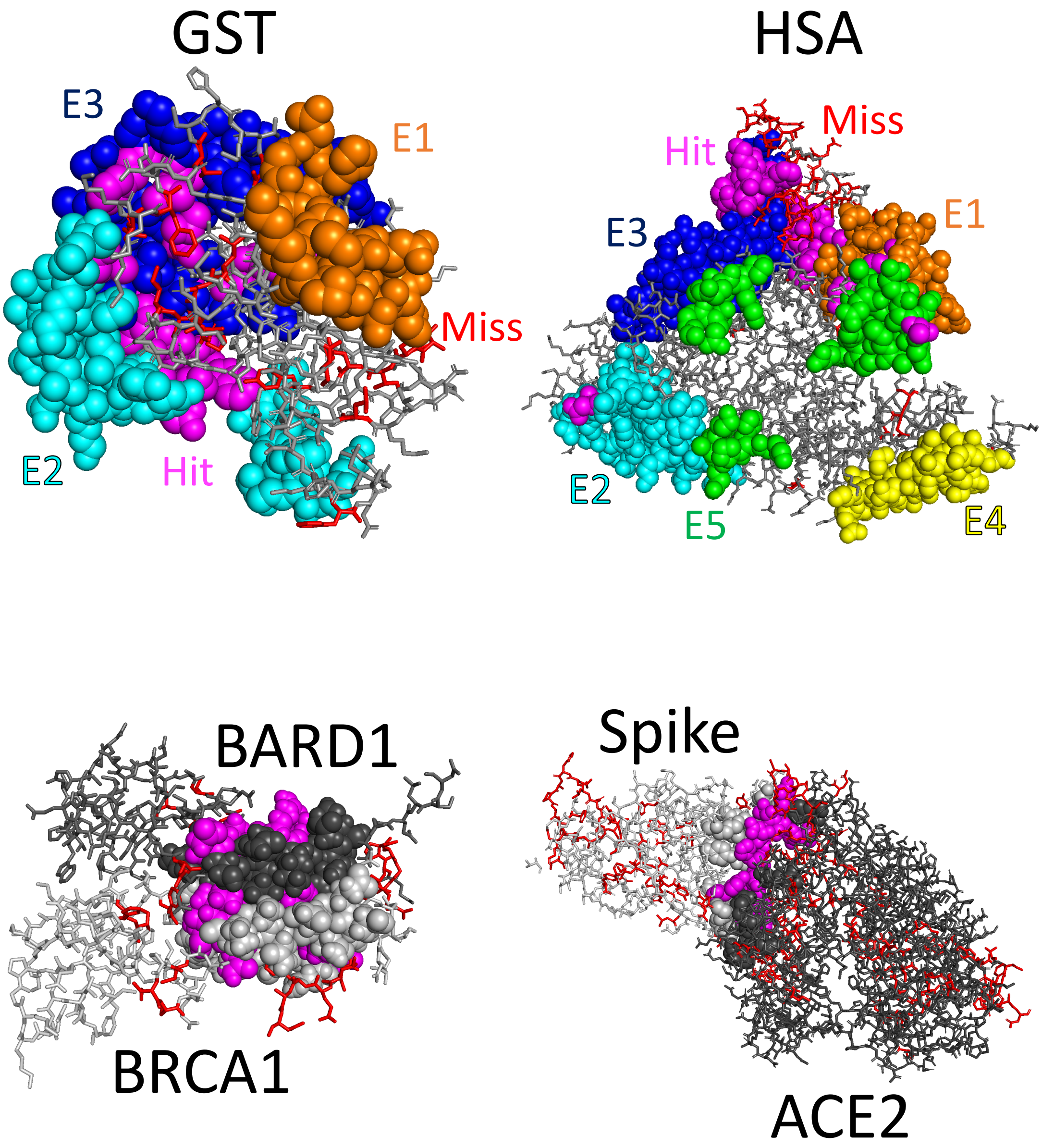


Figure S4: These are the same structures shown in Figure 5, but rotated 180° about the y-axis. With Nb discovery (Top), we highlight overlap between our top scoring residues and epitopes previously validated through cross-linking mass spectrometry. The magenta spheres indicate this overlap. Each other colored sphere indicates a residue within a validated epitope region. The parenthetical percentages indicate the percentage of Nb that bound at each site during the experimental validation. The red sticks correspond to top scoring sites that did not fall in any validated region, and the gray sticks indicate all other residues within the protein sequence. With protein optimization (Bottom), we show structures of each protein bound to its target. The spheres indicate residues known to play a role in protein-target binding. Those shaded magenta were identified by the per-residue scoring. Red sticks indicate residues outside this binding region that were also identified. Dark gray sticks indicate $s_{target}$ residues, and light gray $s_{sequence}$ residues. The structures we used are given by PDB IDs 1DUG, 1AO6, 1JM7, and 6M0J.

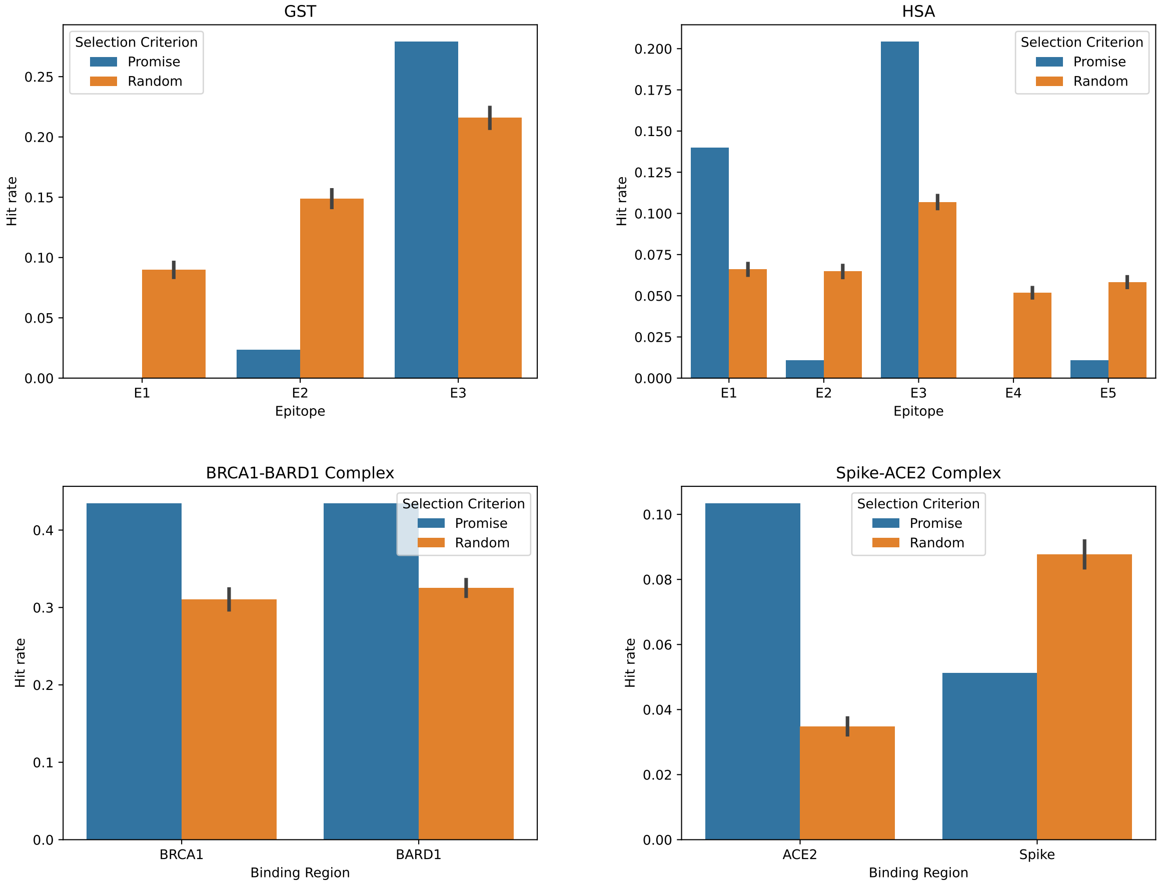


Figure S5: The frequency (ie. "Hit rate") at which PROMISE SCORE selected and randomly selected residues fall within a known binding region for a given protein system. The hit rate obtained via random selection is given as a mean over 50 replicates $\pm$ 1 SEM.

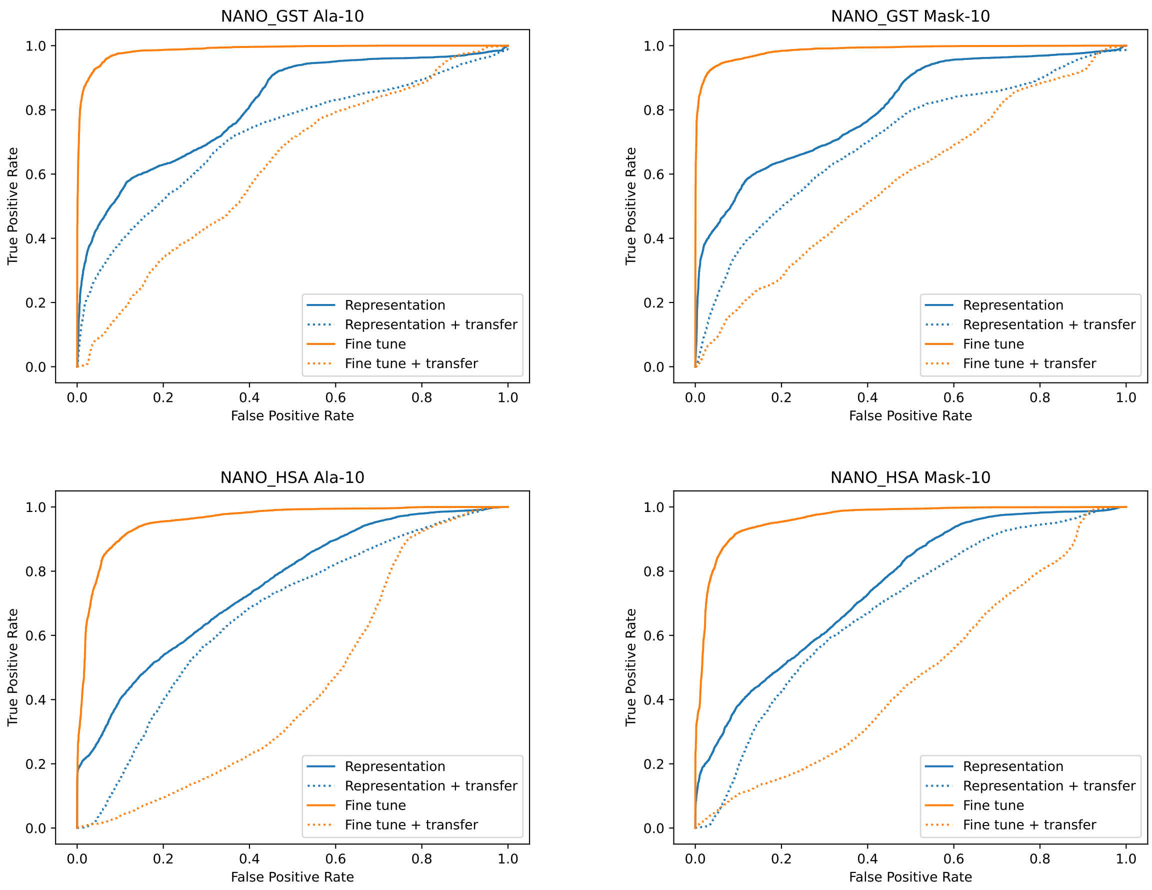


Figure S6: ROC curves for models trained to classify strong and weak sequences with Nb discovery. The top row shows Nano-GST, and the bottom Nano-HSA. The left column shows models trained with a length 10 alanine linker, and the right columns shows models trained with a length ten mask linker. Orange curves correspond to models that used fine-tuning, and blue those that did not. Solid curves correspond to models applied in a traditional machine learning paradigm, whereas dashed lines used transfer learning. Fine-tuned models used in the traditional setting were very strong, regardless of linker type or length.

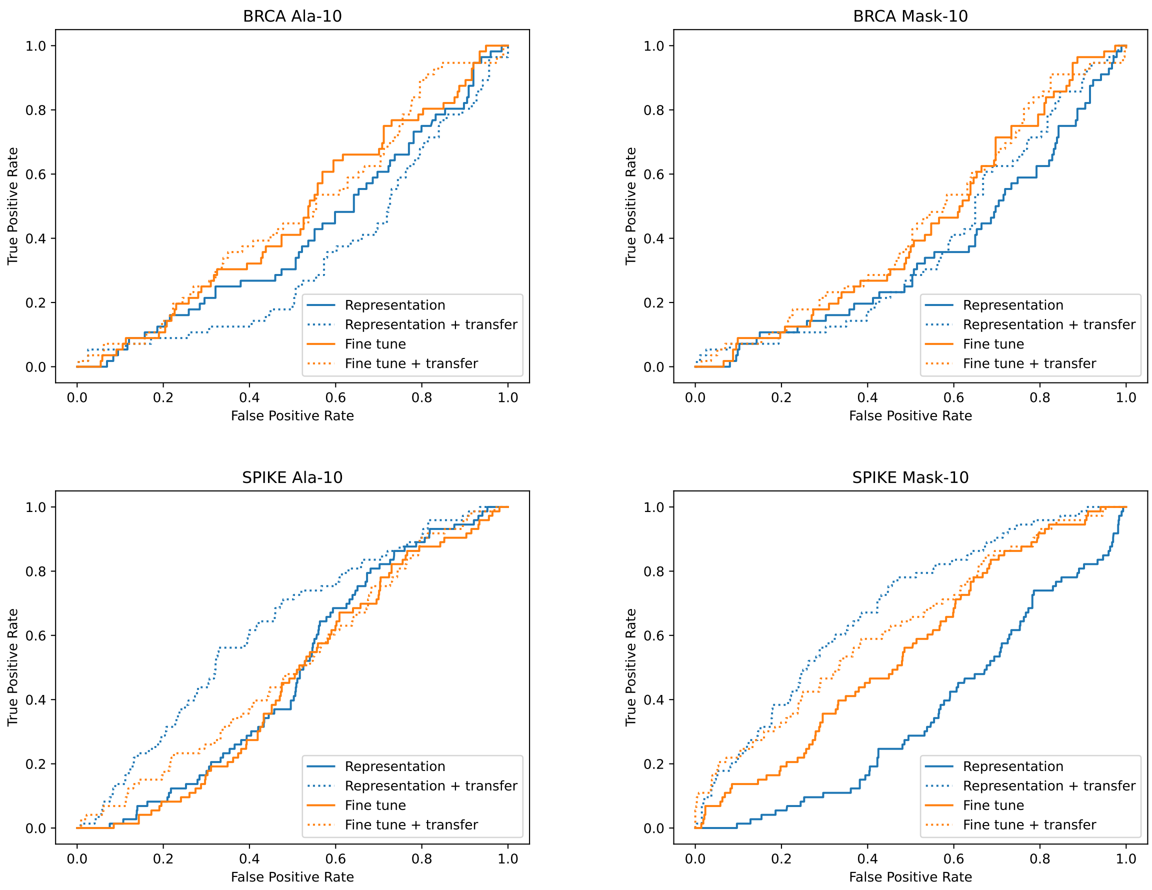


Figure S7: ROC curves for models trained to classify strong and weak sequences in protein optimization. The top row shows BRCA1, and the bottom Spike. The left column shows models trained with a length 10 alanine linker, and the right columns shows models trained with a length ten mask linker. Orange curves correspond to models that used fine-tuning, and blue those that did not. Solid curves correspond to models applied in a traditional machine learning paradigm, whereas dashed lines used transfer learning. While all models learned in protein optimization were of lower quality, we saw evidence that transfer learner led to model improvement.
